# Forest-weighted causal correlation: A scalable framework for causal inference in ecological time series

**DOI:** 10.64898/2026.10.01.756118

**Authors:** Shigeki Kishi

## Abstract

1. Detecting causal relationships from ecological time series remains challenging because community dynamics are often nonlinear, high-dimensional, noisy, and strongly synchronized. These characteristics can reduce the reliability of existing causal inference methods and complicate the identification of ecological interactions in complex natural systems.
2. I introduce Forest-weighted Causal Correlation (FORCC), a machine-learning-based framework that infers directional causality from adaptive neighborhood structures learned by random forests. Rather than evaluating causal direction through cross-map reconstruction, FORCC quantifies directional dependence from the preservation of neighborhood structure using rank-based associations among forest-weight matrices.
3. Simulations of coupled nonlinear systems showed that FORCC accurately recovered causal direction while substantially reducing synchronization-induced false-positive detections relative to conventional convergent cross mapping. Chatterjee’s ξ exhibited the strongest resistance to synchronization-induced errors among the evaluated rank-based statistics.
4. Application to a 12-year marine community time series from Maizuru Bay, Japan, identified asymmetric relationships between individual taxa and community dynamics, revealing candidate driver-like and passenger-like roles that were not readily explained by seasonal synchrony alone.
5. By combining adaptive neighborhood learning with rank-based causal inference, FORCC provides a scalable framework for causal discovery in ecological time series. The method offers a practical tool for screening causal relationships, generating ecological hypotheses, and investigating complex interactions in multivariate natural systems.

## INTRODUCTION

Real-world dynamic systems, from ecological communities to evolutionary and environmental processes, are shaped by complex interactions among many components. Advances in monitoring technologies have generated unprecedented volumes of multivariate time-series data, creating new opportunities to uncover the mechanisms that govern natural systems (Cheng et al., 2015; Misra et al., 2022). Identifying causal relationships from observational time series is therefore a central challenge across ecology and evolution, where understanding how species, populations, and environmental drivers influence one another is essential for predicting ecosystem responses to environmental change (Runge et al., 2019).

Convergent Cross Mapping (CCM) has become one of the most widely used approaches for causal inference in nonlinear dynamical systems (Sugihara et al., 2012). Rooted in Takens’ embedding theory (Takens, 1981), CCM exploits the principle that information about a causal variable becomes embedded in the dynamics of an affected variable. When one variable influences another, local neighborhood relationships in the reconstructed state space of the effect retain information about the cause, enabling directional causality to be inferred from cross-mapping performance (Runge et al., 2023). Applications of CCM have revealed causal relationships in a wide variety of ecological systems, including species interactions, ecosystem processes, and environmental forcing (Sugihara et al., 2012; Ye & Sugihara, 2016).

Despite its broad utility, causal inference in ecological time series remains challenging. Ecological systems are frequently characterized by high dimensionality, observational noise, seasonal forcing, and strong synchronization among interacting variables (Cobey & Baskerville, 2016; Park et al., 2023). Conventional state-space approaches typically quantify neighborhood similarity using Euclidean distances. However, distance measures can become increasingly unreliable in high-dimensional settings because noise and irrelevant variation obscure dynamically meaningful neighbors (Aggarwal et al., 2001; Chang et al., 2021). In addition, strong coupling among variables can induce generalized synchronization, increasing the risk of spurious causal detections and complicating the interpretation of inferred ecological interactions (Mayr et al., 2025; Park et al., 2023). These challenges motivate the development of causal inference methods that can more flexibly and correctly capture local dynamical structure while remaining computationally practical for large ecological datasets.

Machine-learning approaches provide an alternative way to characterize local dynamical neighborhoods. In particular, generalized random forests generate adaptive similarity representations based on repeated partitioning of data into locally homogeneous regions (Athey et al., 2019). The resulting forest weights quantify the degree to which observations share common local structure and can provide a flexible representation of neighborhood relationships that is less dependent on fixed distance metrics (Lin & Jeon, 2006). Rather than relying solely on geometric distances, forest-weight representations allow local relationships to be learned directly from the data, potentially improving the characterization of complex ecological dynamics.

Here, I introduce Forest-weighted Causal Correlation (FORCC), a causal-inference framework that evaluates directional dependence from machine-learning-derived neighborhood structure (Fig. 1). FORCC quantifies the correspondence between localized forest-weight neighborhoods using rank-based association measures, allowing causal direction to be inferred from the preservation of neighborhood structure between variables. I evaluate the performance of FORCC using coupled nonlinear simulations and compare its behavior with existing causal inference approaches. I then apply the framework to a 12-year marine community time series from Maizuru Bay, Japan (Ushio et al., 2018), to examine causal relationships between individual taxa and community dynamics. By combining adaptive neighborhood representations with rank-based causal inference, FORCC provides a scalable framework for causal discovery in ecological time series and offers a practical tool for investigating complex interactions in natural systems.

**FIGURE 1.**
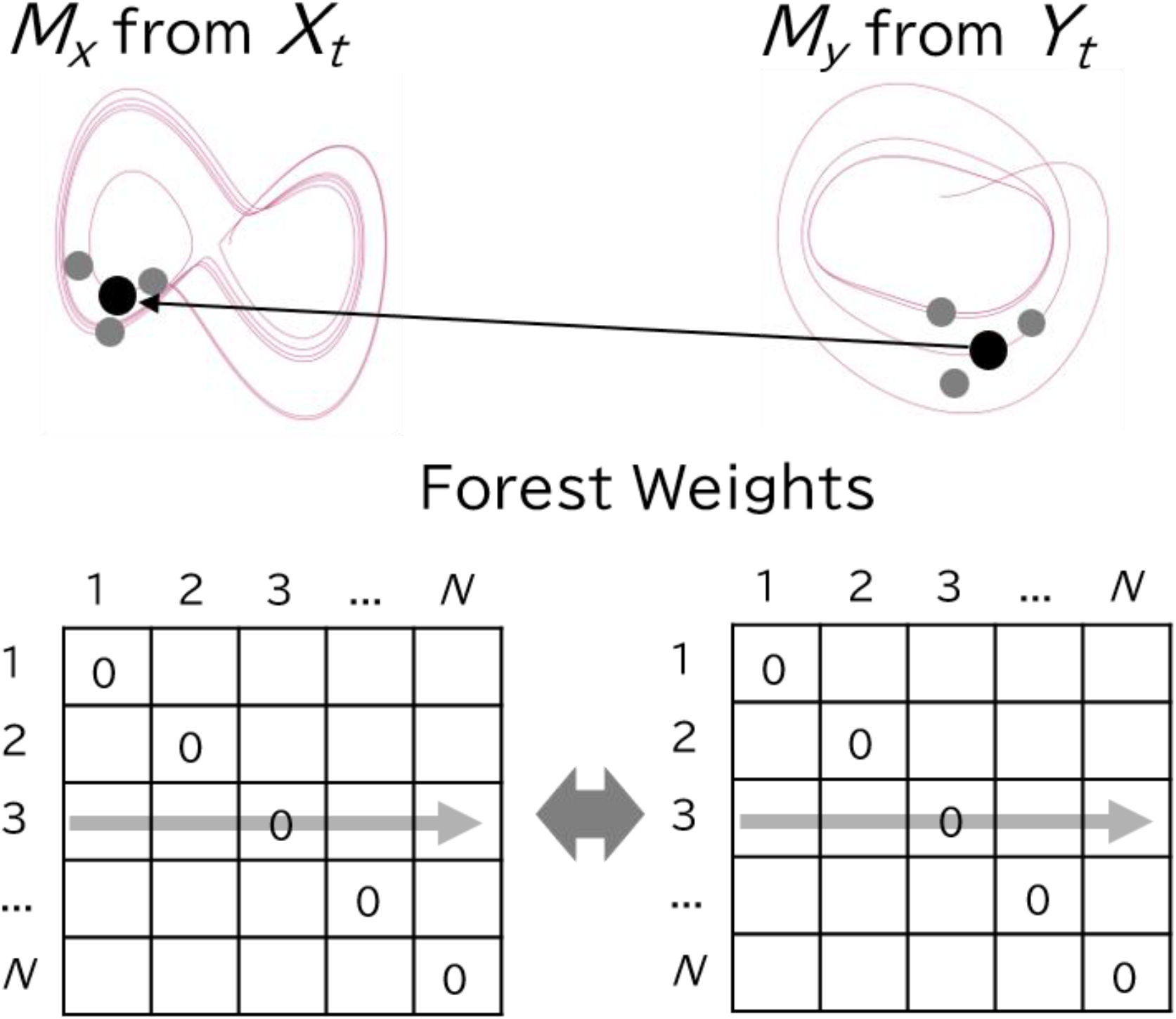
Conceptual framework of Forest-weighted Causal Correlation (FORCC). Under directional causality (*X* → *Y*), local neighborhoods on the effect manifold (*M*_*Y*_) correspond to neighborhoods on the causal manifold (*M*_*X*_). Forest weights derived from generalized random forests encode these neighborhood relationships as *N* × *N* proximity matrices. FORCC detects causality by evaluating rank correlations between corresponding rows of the effect and causal forest-weight matrices, thereby bypassing explicit state-space reconstruction.

## METHODS

### Overview of Forest-weighted Causal Correlation

Forest-weighted Causal Correlation (FORCC) infers directional causal relationships by testing whether local neighborhood structure learned from the dynamics of an affected variable is preserved in the dynamics of a putative causal variable. Unlike conventional cross-mapping approaches, FORCC does not evaluate causality by reconstructing the values of a putative causal variable from the reconstructed states of an affected variable. Instead, it represents local dynamical neighborhoods using forest weights and measures the correspondence between the resulting neighborhood structures with rank-based dependence statistics.

The FORCC workflow consists of five principal steps (Fig. 1). First, delay-coordinate vectors are constructed from each time series to represent its local dynamical states. Second, a regression forest is fitted to predict the forward dynamics of each variable from its delay coordinates. Third, the fitted forest is used to construct a sample-weight matrix describing the adaptive neighborhood surrounding each state. Fourth, corresponding rows of the weight matrices for the putative cause and effect are aligned at the specified causal lag and compared using rank-based dependence measures. Finally, the mean dependence across local states is evaluated against a null distribution generated from phase-randomized surrogate time series.

Hereafter, I use *X* → *Y* to denote the hypothesis that variable *X* causally influences variable *Y*. Under this hypothesis, the local dynamical neighborhoods identified from the states of *Y* are expected to retain information about the corresponding states of *X*. Consequently, neighborhood relationships encoded in the forest-weight matrix of *Y* should be more strongly preserved in the forest-weight matrix of *X* than expected under a null model in which directional dependence has been disrupted.

### Construction of dynamical state vectors

For a univariate time series *y*_*t*_, the dynamical state at time *t* was represented by an *E*-dimensional delay-coordinate vector,

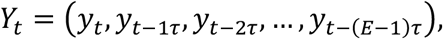

where *E* is the embedding dimension and *τ* is the delay interval. I used *τ* = 1, corresponding to one sampling interval, throughout the analyses.

The embedding dimension was selected independently for each variable by evaluating out-of-sample prediction of its forward dynamics. Candidate embedding dimensions from *E* = 1 to *E* = 12 were evaluated for the Maizuru Bay community analysis. For the simulation analyses, *E* = 2 was used because preliminary optimization identified this value as suitable for the coupled logistic-map system. The dimension producing the highest out-of-sample Pearson correlation between observed and predicted values was selected as the optimal dimension, *E*^∗^.

To reduce information leakage arising from temporal dependence, prediction skill was evaluated using out-of-bag prediction. Ties in prediction skill were resolved by selecting the smallest dimension attaining the maximum skill.

For a multivariate community containing *K* taxa, the state vector at time *t* was constructed by combining the lagged coordinates of the component time series,

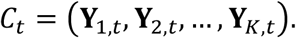

For each focal taxon *s* in the empirical analysis, a separate community state vector *C*_−*s,t*_ was constructed from the remaining 14 taxa. Thus, the definition of the community and its corresponding forest-weight matrix differed among focal taxa.

### Estimation of forest-weight neighborhoods

Local dynamical neighborhoods were represented using sample weights derived from generalized random forests. For a univariate time series, a regression forest was fitted to predict the forward value *y*_*t*+1_ from the corresponding state vector **Y**_*t*_. Each forest contained 500 trees. This number was chosen because preliminary analyses indicated that forest-weight estimates became stable at approximately this forest size, with little additional improvement from larger ensembles.

A decision tree recursively partitions the predictor space into terminal nodes containing observations with similar predictor-response relationships. Observations that repeatedly occur in the same terminal nodes across trees can therefore be treated as belonging to similar local dynamical neighborhoods. For a query state **Y**_*i*_, the weight assigned to observation **Y**_*j*_ was calculated as

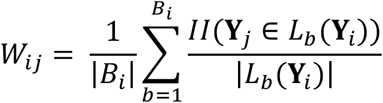

where *b* indexes individual trees in the forest and *B*_*i*_ denotes the number of trees contributing to the weight calculation for query observation *i, L*_*b*_(**Y**_*i*_) is the terminal node containing **Y**_*i*_ in tree b, ∣*L*_*b*_(**Y**_*i*_)∣ is the number of training observations in that node, and II[⋅] is the indicator function.

The resulting *N*×*N* matrix,

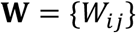

provides an adaptive representation of local neighborhood structure. Each row contains the weights assigned to all observations relative to one query state. Large positive weights indicate observations that were frequently assigned to the same terminal nodes as the query state, after accounting for terminal-node size.

To prevent a query observation from being treated as its own neighbor, the diagonal element *W*_*ii*_ was set to zero before calculating the causal statistics. Only observations with positive weights in the effect-variable matrix were retained as local neighbors. Forest models were fitted using the grf package. I used the default settings unless referring to the hyper-parameter values (honesty = TRUE, honesty.fraction = 0.5, sample.fraction = 0.5, mtry = min(ceiling(sqrt(ncol(x))+20), ncol(x)), min.node.size = 5, alpha = 0.5, imbalance.penalty = 0).

For the multivariate community analysis, a multivariate regression forest was fitted to predict the forward abundance vector of all community members simultaneously from the corresponding community state vector. Tree splits were selected using the default multivariate splitting criterion implemented in multi_regression_forest().

### Directional dependence between forest-weight matrices

To test the hypothesis *X* → *Y*, I treated *Y* as the putative effect and *X* as the putative cause. For each query state *i*, I first identified the observations receiving positive weights in the effect matrix,

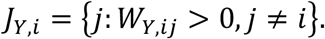

The corresponding weight vectors were then extracted from the effect and cause matrices,

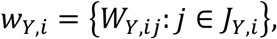

and

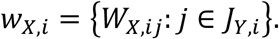

Thus, neighborhoods were defined from the putative effect, and the preservation of those same neighborhood relationships was evaluated in the putative cause. Reversing the roles of the matrices provided the statistic for the opposite causal hypothesis, *Y* → *X*.

A query state was retained only when the effect matrix contained at least five positive-weight neighbors after temporal alignment. This minimum was imposed to avoid calculating rank-based statistics from extremely small local samples. Because the choice of this threshold may influence the results, I evaluated its sensitivity using minimum neighborhood sizes of 5, 10, and 20 in a supplementary analysis.

The cause and effect matrices were aligned according to their prediction-target times. For the causal hypothesis *X*_*t*_→*Y*_*t*+1_, the neighborhood structure associated with the state of *Y* at time *t* was compared with the corresponding neighborhood structure of *X* at time *t*. This alignment ensures that the two forest-weight rows refer to states associated with the same dynamical transition.

Because delay-coordinate embedding represents each state using multiple historical observations, the temporal reference point of an embedded state differs from that of the original time series. To align neighborhood structures associated with the same underlying dynamical event, the effective lag used in FORCC was defined as

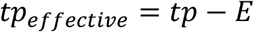

where *tp* is the causal lag and *E* is the embedding dimension. This adjustment compensates for the temporal displacement introduced by the embedded representation and aligns rows corresponding to the same dynamical transition.

### Rank-based causal statistics

Local correspondence between the effect and cause weight vectors was quantified using Spearman’s rank correlation coefficient, Kendall’s rank correlation coefficient, and Chatterjee’s rank correlation coefficient. Spearman’s *ρ* and Kendall’s *τ* measure monotonic rank association, whereas Chatterjee’s *ξ* can detect more general functional dependence that need not be monotonic.

For Chatterjee’s coefficient, the paired observations were ordered according to the ranks of the effect weights. Let *M*_*i*_=∣*J*_*Y,i*_∣ be the number of neighbors retained for query state *i*, and let *R*_*X,i*,(*j*)_ denote the rank of the cause weight paired with the *j*th ordered effect weight. In the absence of ties, the coefficient was calculated as

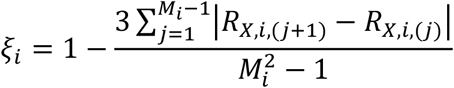

Chatterjee’s *ξ* is directional. I therefore treated the effect-weight ranks as the ordering variable and the corresponding cause-weight ranks as the dependent variable.

Chatterjee’s ξ was calculated using the implementation provided in the XICOR package (version 0.4.0), which was also used to handle ties.

A single statistic for a causal hypothesis was obtained by averaging the local statistics across all valid query states,

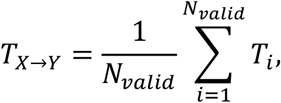

where *T*_*i*_ is the local rank-dependence statistic and *N*_*valid*_ is the number of query states meeting the neighborhood-size requirement.

Higher values of Spearman’s *ρ*, Kendall’s *τ*, or Chatterjee’s *ξ* indicate stronger preservation of the effect-defined neighborhoods in the putative cause. Directional causality was inferred only when an observed statistic exceeded the corresponding upper bound of its surrogate null distribution. To evaluate distributional divergence between row-normalized positive forest weights, Kullback–Leibler (KL) divergence was computed as

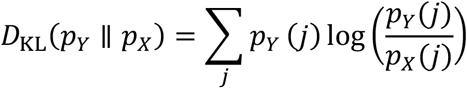

where small epsilon values were added to avoid zero division.

### Surrogate testing

The statistical significance of each directional FORCC statistic was evaluated using phase-randomized surrogate time series generated with the Ebisuzaki method (Ebisuzaki, 1997). Phase randomization preserves the approximate power spectrum and linear autocorrelation structure of the original series while disrupting its phase relationships with other variables.

For each causal hypothesis, *S* independent surrogate realizations of the putative cause time series were generated. The full FORCC procedure, including the construction of state vectors, fitting of regression forests, extraction of forest weights, temporal alignment, and calculation of the mean rank-dependence statistic, was repeated for each surrogate.

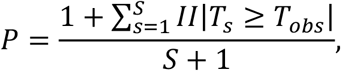

where *T*_obs_ is the statistic calculated from the observed time series and *T*_*s*_ is the statistic from surrogate realization *s*. The addition of one to the numerator and denominator prevents zero-valued empirical *P*-values.

I used 200 surrogate realizations for the detailed illustration of a single simulated dataset and 100 surrogate realizations for each replicate in the simulation sensitivity analysis and empirical community analysis. A causal relationship was considered statistically supported when the one-sided empirical *P*-value was less than 0.05.

### Evaluation using coupled nonlinear dynamics

I evaluated the ability of FORCC to identify directional causality using a unidirectionally coupled logistic-map system,

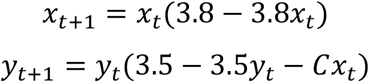

where *C* determines the strength of the causal influence from *X* to *Y*. The system contains a known causal link *X*→*Y* and no direct causal link *Y*→*X*.

Initial values were sampled independently from a uniform distribution between 0.1 and 0.9. Each simulation was run for a burn-in period of 100 time steps, which was discarded before analysis.

Independent Gaussian measurement noise was added to each simulated trajectory,

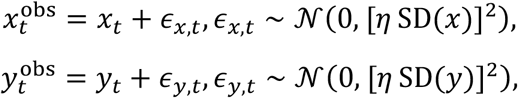

where *η* = 0.01. Thus, the standard deviation of the added measurement noise was 1% of the standard deviation of the corresponding noise-free trajectory.

### Illustration of local neighborhood correspondence

To illustrate the directional preservation of neighborhood structure, I generated one representative dataset with *C* = 0.1, *N* = 300, and *η* = 0.01. Fifty query states were randomly selected for visualization. For each query state, positive weights from the effect matrix and the corresponding elements from the cause matrix were extracted for both causal directions. The same comparisons were repeated after converting weights to within-row ranks.

Kernel density estimates were used to summarize the distributions of local Spearman’s *ρ*, Kendall’s *τ*, and Chatterjee’s *ξ* across all valid query states. These local distributions were used only to visualize directional differences. Statistical inference was based on the cross-state mean statistics and their surrogate distributions.

### Performance across coupling strengths

Sensitivity to coupling strength was evaluated at

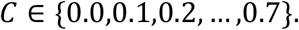

For each coupling strength, 200 independently initialized datasets were generated. Each dataset was tested in both the true causal direction, *X*→*Y*, and the non-causal direction, *Y*→*X*, using 100 phase-randomized surrogates.

The true-positive rate was calculated as the proportion of the 200 trials in which the null hypothesis was rejected for *X*→*Y*. The false-positive rate was calculated as the proportion in which the null hypothesis was rejected for *Y*→*X*. At *C* = 0, rejection in either direction was interpreted as a false-positive result.

### Application to the Maizuru Bay marine community

I evaluated the applicability of FORCC to a 12-year observational time series collected from a coastal marine community in Maizuru Bay, Japan. The dataset contains biweekly abundance observations for 15 dominant marine taxa, together with measurements of surface and bottom water temperature. For community analyses, multivariate responses were modeled using multi_regression_forest().

All abundance and temperature series were standardized separately to zero mean and unit variance before analysis. Standardization prevented differences in measurement scale and abundance variability from dominating the forest-fitting procedure.

For each focal taxon *s*, the remaining 14 taxa were combined to create a focal-taxon-specific community matrix *C*_−*s*_. Directional dependence was tested for both

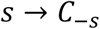

and

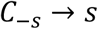

A separate community forest-weight matrix was therefore constructed for each focal taxon. This leave-one-taxon-out design prevented the focal taxon from contributing directly to the community state against which it was tested.

The embedding dimension was selected independently for each variable by evaluating out-of-sample prediction of its forward dynamics. Candidate dimensions from *E* = 1 to *E* = 12 were compared by fitting regression forests that predicted *y*_*t*+1_ from **Y**_t_. The upper limit of 12 was chosen to allow the embedded state vectors to capture the dominant annual seasonal cycle present in the Maizuru Bay time series. The dimension producing the highest out-of-sample Pearson correlation between observed and predicted values was selected as the optimal dimension, *E*^∗^. The optimal embedding dimensions selected by out-of-sample prediction were *E*^∗^ = 11 for the community state, *E*^∗^ = 11 for surface water temperature, and *E*^∗^ = 9 for bottom water temperature.

For each focal taxon, significance was evaluated using 100 phase-randomized surrogate time series generated with the Ebisuzaki method. The focal taxon was replaced by surrogate realizations while the observed dynamics of the remaining 14-taxon community were retained unchanged. Consequently, the surrogate analyses tested whether observed taxon-community relationships exceeded expectations generated by the observed community background alone. This procedure preserved the observed community structure, including seasonal variation and interspecific synchrony, while disrupting the temporal structure of the focal taxon.

Observed FORCC statistics were compared with the distribution obtained from the surrogate focal-taxon series. Relationships exceeding surrogate expectations were interpreted as directional associations that could not be readily explained by the preserved community dynamics alone.

### Computational environment and reproducibility

All simulations and statistical analyses were conducted in R version 4.4.0 (R Core Team, 2025). Generalized random forests and forest-weight extraction were implemented using the grf package, version 2.6.1 (Tibshirani et al., 2026). Conventional CCM was implemented using rEDM, version 0.7.5 (https://github.com/ha0ye/rEDM). Chatterjee’s *ξ* was calculated using XICOR, version 0.4.0. MDR S-map and multiview distance matrices were calculated using rUIC, version 0.9.15 (https://github.com/yutakaos/rUIC), and macam, version 0.2.5 (https://github.com/ong8181/macam).

Computationally intensive simulation and surrogate analyses were conducted on the SHIHOU high-performance computing system at the National Agriculture and Food Research Organization. Analyses were parallelized using the furrr package with a multisession backend.

## RESULTS

### Local neighborhood structure preserves directional information

To examine whether directional causality is reflected in local neighborhood structure, I first compared corresponding rows of forest-weight matrices derived from coupled nonlinear dynamics (Fig. 2). Under the true causal direction (*X*→*Y*), forest-weight neighborhoods identified from the effect variable showed stronger correspondence with those of the putative cause than under the non-causal direction (*Y*→*X*) (Fig. 2). This pattern was evident both for the raw forest weights and for their within-row ranks, indicating that neighborhood relationships identified from the effect variable were more consistently preserved in the causal variable than in the reverse direction.

**FIGURE 2.**
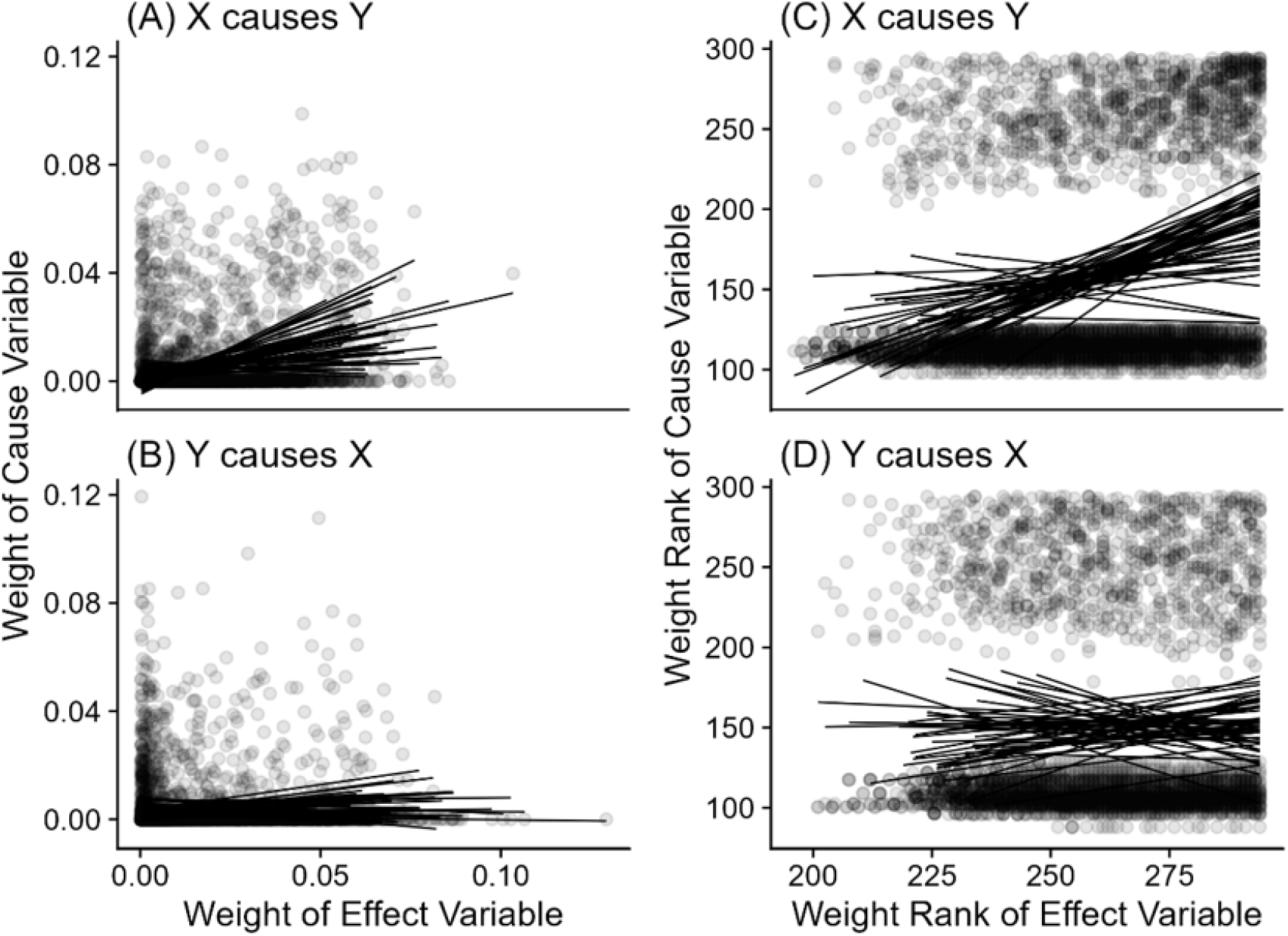
Directional asymmetry of forest weights and their ranks. (A, B) Relationships between forest weights from the effect manifold and the corresponding weights from the causal manifold for the causal (*X* → *Y*) and non-causal (*Y* → *X*) directions, respectively. (C, D) Corresponding relationships after conversion to within-row ranks. Gray points indicate neighborhood relationships extracted from 50 randomly selected query states, and black lines represent query-specific least-squares trend lines included for visualization only. Correspondence between effect-defined and cause-defined neighborhoods is stronger in the causal direction (*X* → *Y*) than in the non-causal direction (*Y* → *X*).

Across all valid query states, the distributions of Spearman’s *ρ*, Kendall’s *τ*, and Chatterjee’s *ξ* were shifted toward larger values in the true causal direction relative to the non-causal direction (Fig. 3A-C). This pattern indicates that local neighborhoods identified from an affected variable retain information about the structure of the causal variable, supporting the central premise of FORCC. A similar pattern was observed for KL divergence, which was lower in the causal direction, indicating greater similarity between the underlying forest-weight distributions (Fig. 3D).

**FIGURE 3.**
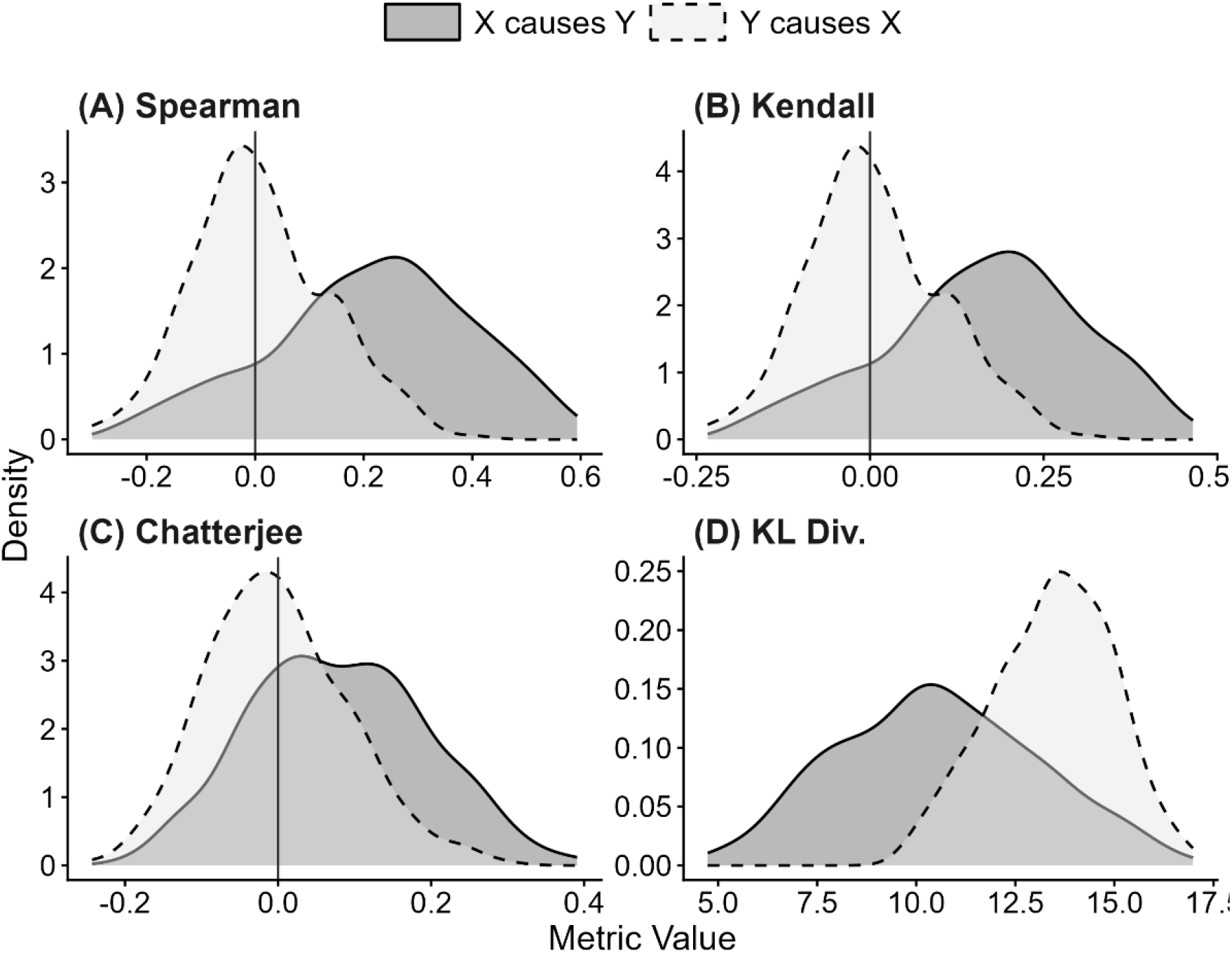
Distributions of topological similarity metrics under causal and non-causal directions. Kernel density estimates of row-wise metric values calculated from forest-weight matrices for the causal direction (*X* → *Y*) and the non-causal direction (*Y* → *X*). Metrics include (A) Spearman’s *ρ*, (B) Kendall’s *τ*, (C) Chatterjee’s *ξ*, and (D) Kullback-Leibler (KL) divergence. Rank-correlation metrics show higher values in the causal direction, whereas KL divergence is lower, consistent with stronger topological similarity between corresponding neighborhood structures.

### FORCC accurately identifies causal direction

The observed rank-dependence statistics differed strongly from expectations generated by phase-randomized surrogate time series (Fig. 4). For the true causal direction (*X*→*Y*), the observed values of Spearman’s *ρ* and Chatterjee’s *ξ* exceeded the upper bounds of the surrogate distributions. For KL divergence, support for causality was defined only when the observed value fell below the lower 95% surrogate bound. Accordingly, X→Y exhibited the expected reduction in KL divergence relative to surrogate expectations, whereas the elevated KL divergence observed for Y→X was interpreted as evidence of reduced neighborhood similarity rather than evidence of causality.

**FIGURE 4.**
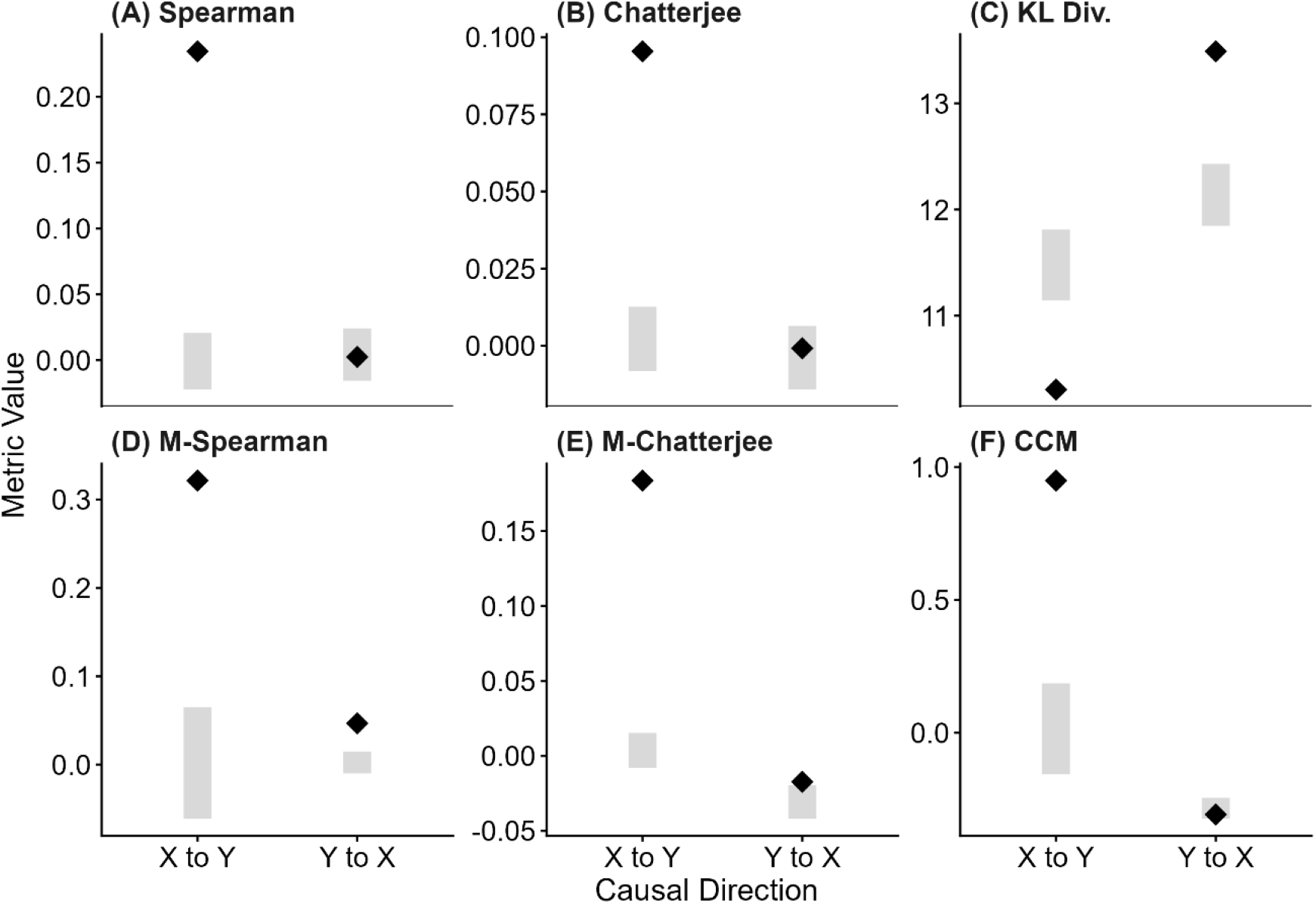
Statistical evaluation of causal directions using phase-randomized surrogate data. Black diamonds indicate observed mean metric values and gray bars show 95% confidence intervals obtained from phase-randomized surrogate time series. (A, B) Spearman’s *ρ* and Chatterjee’s *ξ* calculated from forest-weight matrices. (C) KL divergence calculated from forest-weight matrices. (D, E) Corresponding rank- correlation metrics calculated from multiview distance matrices (MVD). (F) Conventional convergent cross mapping (CCM). Evidence for causality was defined as an observed value exceeding the upper 95% surrogate bound for rank-based statistics or falling below the lower 95% surrogate bound for KL divergence. Observed KL-divergence values exceeding the upper surrogate bound were not interpreted as evidence of causality.

Applying the same rank-based framework to multiview distance matrices also detected the true causal direction, but discrimination between causal and non-causal directions was weaker than for forest-weight matrices. Conventional convergent cross mapping (CCM) correctly recovered the known causal direction in this representative dataset. Together, these comparisons suggest that adaptive forest-weight neighborhoods provide additional information that improves the separation of causal and non-causal relationships.

### Robustness to generalized synchronization

I next evaluated FORCC across a gradient of coupling strengths (*C* = 0.0−0.7) to determine whether the method remained reliable under increasing synchronization between variables (Fig. 5). All evaluated methods detected the true causal direction (*X*→*Y*) with high frequency once coupling was present. Detection rates approached 100% for all FORCC metrics and for CCM at moderate to strong coupling strengths.

**FIGURE 5.**
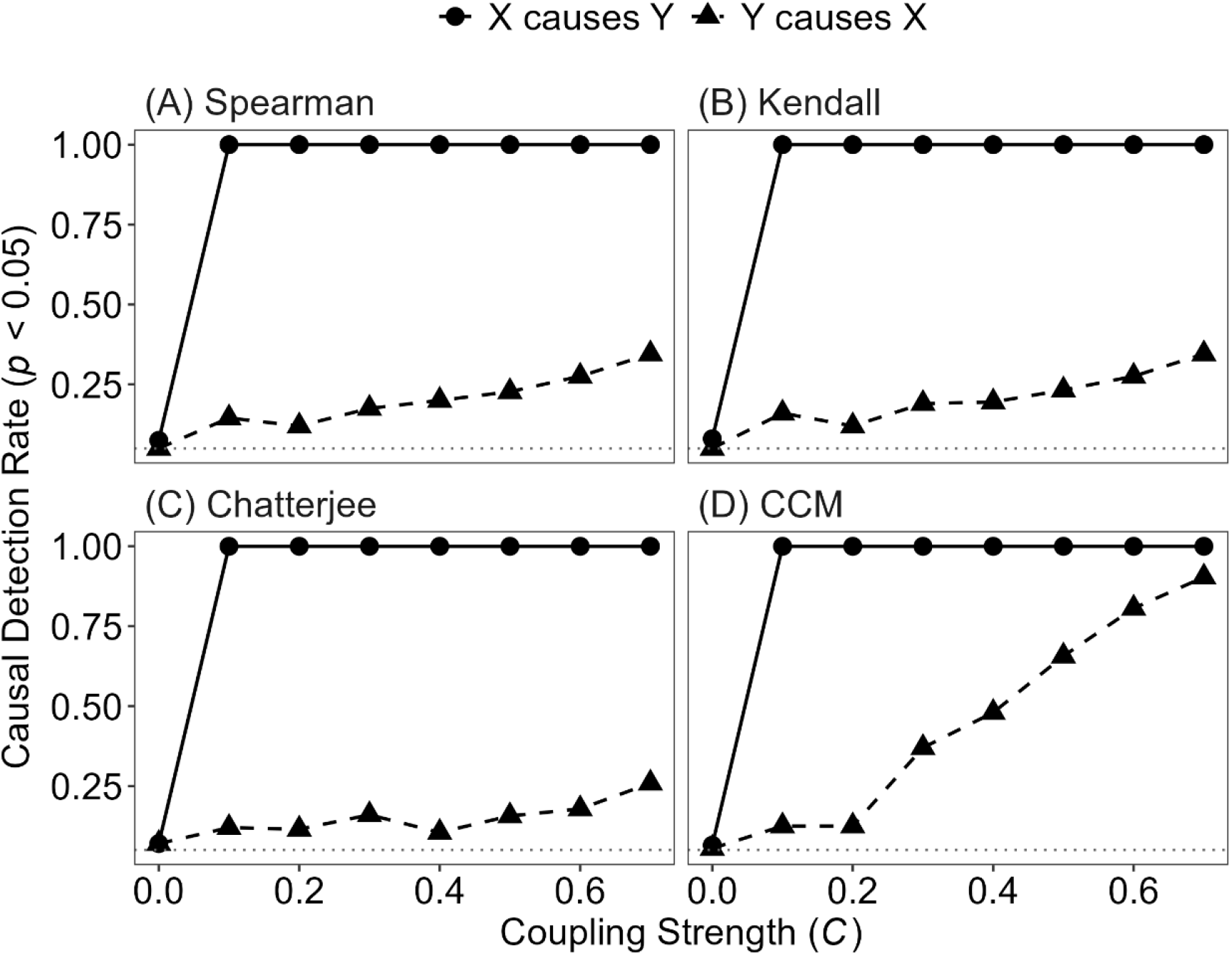
Causal detection rates across coupling strengths. Detection rates (*P* < 0.05) obtained from 200 independent simulations of a unidirectionally coupled logistic map. (A-C) FORCC using Spearman’s *ρ*, Kendall’s *τ*, and Chatterjee’s *ξ*, respectively. (D) Convergent cross mapping (CCM). Circles denote detection rates for the true causal direction (*X* → *Y*), whereas triangles denote false-positive rates for the non-causal direction (*Y* → *X*). The dotted horizontal line indicates the nominal significance level (*α* = 0.05). All methods recovered the causal direction under non-zero coupling, whereas FORCC maintained lower false-positive rates than CCM under stronger coupling.

More substantial differences emerged for the non-causal direction (*Y*→*X*). False-positive detections increased markedly for CCM as coupling strength increased, reaching 90.4% at the strongest coupling level examined. In contrast, all FORCC variants produced substantially fewer false positives than CCM. Chatterjee’s *ξ* showed the greatest resistance to synchronization-induced errors, maintaining false-positive rates of 10.5%-25.9% across the strongest coupling regimes. Although false positives were not eliminated completely, these results demonstrate that FORCC substantially mitigates the influence of generalized synchronization on causal inference.

### Identifying asymmetric relationships in a marine community

To evaluate the applicability of FORCC to ecological time series, I analyzed a 12-year observational record of a coastal marine community from Maizuru Bay, Japan (Ushio et al., 2018). FORCC was applied to test directional relationships between individual taxa and the remainder of the community while accounting for the temporal structure of focal taxa through phase-randomized surrogate time series.

Several taxa, including Japanese jack mackerel (*Trachurus japonicus*), largescale blackfish (*Girella punctata*), and spotty-belly greenling (*Halichoeres poecilopterus*), exhibited directional associations that exceeded surrogate expectations (Fig. 6A), suggesting that their dynamics were linked to community-level structure in a manner not readily explained by shared seasonal variation alone.

**FIGURE 6.**
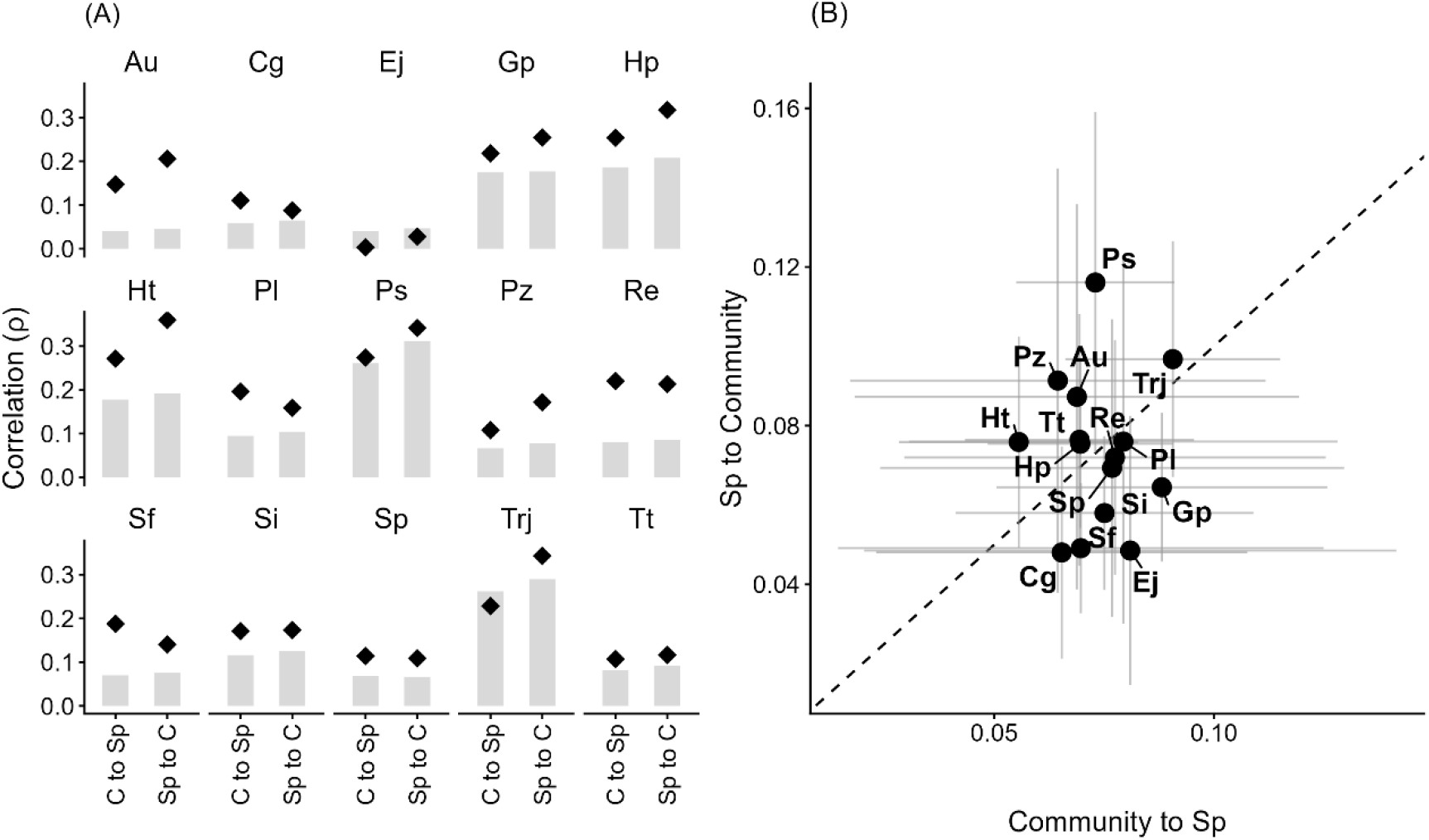
Empirical application to a 12-year marine community time series from Maizuru Bay. (A) Bidirectional causal inference between individual species (Sp) and the remaining 14-species community (C) using FORCC. Black diamonds indicate observed mean Spearman’s *ρ* values and gray bars indicate 95% confidence intervals obtained from phase-randomized surrogate data. Species were considered significant when observed values exceeded the upper 95% surrogate bound. (B) Asymmetric ecological interaction structure inferred using Forest-weighted S-map (FORWS). The horizontal axis shows interaction strength from the community to each species and the vertical axis shows interaction strength from each species to the community. The dashed line indicates equal influence in both directions. Species located above the line exert stronger community-level influence, whereas those below the line are more strongly influenced by community dynamics. Abbreviations: Au, *Aurelia* sp.; Cg, *Chaenogobius gulosus*; Ej, *Engraulis japonicus* (Japanese anchovy); Gp, *Girella punctata*; Hp, *Halichoeres poecilopterus*; Ht, *Halichoeres tenuispinnis*; Pl, *Plotosus lineatus*; Ps, *Pseudolabrus sieboldi*; Pz, *Pterogobius zonoleucus*; Re, *Rudarius ercodes*; Si, *Sebastes inermis*; Sf, *Siganus fuscescens*; Sp, *Sphyraena pinguis*; Trj, *Trachurus japonicus* (Japanese jack mackerel); Tt, *Tridentiger trigonocephalus*.

To further interpret these asymmetric relationships, I quantified directional interaction strengths using Forest-weighted S-map (FORWS). Comparisons of species-to-community and community-to-species influences revealed substantial variation among taxa (Fig. 6B). Japanese jack mackerel (*T. japonicus*) and *Pseudolabrus sieboldi* exhibited stronger inferred influences on community dynamics than they received from the community, whereas Japanese anchovy (*Engraulis japonicus*) and yellowfin goby (*Chaenogobius gulosus*) showed the opposite pattern. These results illustrate how FORCC can help identify candidate driver-like and passenger-like components within complex ecological communities while generating testable hypotheses about community organization.

## DISCUSSION

This study introduces Forest-weighted Causal Correlation (FORCC), a framework that combines machine-learning-derived neighborhood representations with rank-based measures of directional dependence. In coupled nonlinear simulations, FORCC recovered the known causal direction while producing substantially fewer synchronization-induced false positives than conventional Convergent Cross Mapping (CCM). Applying the framework to a 12-year marine community time series further revealed asymmetric associations between individual taxa and community dynamics. Together, these findings support the central premise of FORCC: local neighborhood relationships learned from an affected variable retain directional information about a putative cause, and this information can be detected without reconstructing the values of the causal variable through cross mapping. FORCC therefore provides a new way to screen causal relationships in complex ecological time series.

The foundation of FORCC is closely related to the principle underlying state-space approaches to causal inference. If *X* causally influences *Y*, information about *X* becomes encoded in the dynamics of *Y*, allowing states of the putative cause to be reconstructed from the state-space representation of the effect (Sugihara et al., 2012; Takens, 1981). Conventional CCM evaluates this correspondence through cross-map reconstruction skill. FORCC instead asks whether local neighborhoods identified from the dynamics of the effect are preserved in the corresponding dynamics of the cause. By converting this question into a comparison between forest-weight matrices, FORCC evaluates directional dependence directly from learned neighborhood relationships. The simulation results show that these neighborhood relationships contain sufficient information to distinguish the true causal direction from the non-causal direction, supporting their use as an alternative basis for causal inference.

The forest-weight representation appears to be an important component of this performance. Conventional state-space methods commonly define local neighborhoods using Euclidean distances, but such distances can be distorted by irrelevant variation, observational noise, and increasing dimensionality (Aggarwal et al., 2001; Chang et al., 2021; Ye & Sugihara, 2016). Forest weights instead describe similarity through repeated co-occurrence of observations within terminal nodes of regression trees (Athey et al., 2019). Because the underlying partitions are learned from the relationship between current states and their forward dynamics, the resulting neighborhoods emphasize aspects of the state space that are informative about dynamical transitions. In the present comparison, applying the same rank-based statistics to multiview distance matrices detected the true causal direction but provided weaker separation between causal and non-causal relationships. This result suggests that the performance of FORCC arises from the combination of two elements: adaptive neighborhood learning through forest weights and direct comparison of neighborhood structure through rank-based dependence.

The differences among the evaluated rank statistics provide further insight into how directional information is represented in forest-weight neighborhoods. Spearman’s *ρ* and Kendall’s *τ* measure monotonic rank association, whereas Chatterjee’s *ξ* can detect broader forms of functional dependence that need not be monotonic (Chatterjee, 2021; Wang et al., 2024). Relationships between corresponding neighborhoods may become non-monotonic when nonlinear attractors are folded, stretched, or locally distorted (Chicharro & Andrzejak, 2009). Consistent with this possibility, Chatterjee’s *ξ* produced the lowest false-positive rates among the evaluated FORCC statistics. This finding does not establish that *ξ* will be optimal for all dynamical systems, but it indicates that statistics capable of detecting non-monotonic dependence may be particularly useful when comparing learned neighborhood structures. The lower KL-divergence values observed in the causal direction further suggest that directional dependence is reflected not only in rank ordering but also in broader similarities between forest-weight distributions. Further theoretical and simulation studies will be needed to determine how the geometry of a dynamical system, the distribution of forest weights, and the number of available local neighbors affect the behavior of these statistics.

A particularly important result for ecological applications was the relative robustness of FORCC to generalized synchronization. Strong coupling can cause the trajectories of interacting variables to become synchronized, making it difficult to distinguish genuine directional influence from the similarity produced by their shared dynamics (Park et al., 2023; Rulkov et al., 1995). In the coupled logistic-map simulations, CCM increasingly detected the non-causal direction as coupling strength increased, whereas FORCC produced substantially lower false-positive rates. Chatterjee’s *ξ* was the most robust statistic, although its false-positive rate still exceeded the nominal significance level under strong coupling. FORCC therefore mitigated, but did not eliminate, synchronization-induced errors. This distinction is important because synchronization is common in ecological time series, where species may respond simultaneously to seasonal forcing, shared environmental conditions, or strongly coupled community processes. A method that reduces false positives under these conditions can provide more reliable candidates for subsequent ecological investigation, but inferred relationships should still be evaluated against alternative explanations and independent biological knowledge.

The Maizuru Bay analysis illustrates how FORCC can be used to investigate directional organization in an observational ecological community. Several taxa, including Japanese jack mackerel (*Trachurus japonicus*), largescale blackfish (*Girella punctata*), and spotty-belly greenling (*Halichoeres poecilopterus*), exhibited directional associations with the remaining community that exceeded phase-randomized surrogate expectations. These results indicate that their relationships with community dynamics were not readily explained by the univariate temporal structures preserved by the surrogate procedure. The complementary analysis of directional interaction strengths further suggested substantial differences among taxa. Japanese jack mackerel (*Trachurus japonicus*) and *Pseudolabrus sieboldi* showed stronger inferred taxon-to-community than community-to-taxon influences, consistent with candidate driver-like roles. Japanese anchovy (*Engraulis japonicus*) and yellowfin goby (*Chaenogobius gulosus*) showed the opposite asymmetry, consistent with candidate passenger-like roles. These classifications should be interpreted as hypotheses about functional organization rather than definitive assignments of ecological roles. Nevertheless, they demonstrate how FORCC can transform multivariate monitoring data into specific, biologically interpretable hypotheses that can be examined using experiments, independent observations, or mechanistic models.

More broadly, FORCC may be useful as a screening tool for ecological causal discovery. Ecological monitoring programs increasingly generate multivariate time series containing many taxa, environmental variables, and sampling occasions. Exhaustively evaluating potential directional relationships in such datasets can be computationally demanding, and the resulting networks can be difficult to interpret when synchronization or shared temporal structure generates numerous apparent associations. FORCC addresses this challenge by summarizing each local dynamical neighborhood as a row of a forest-weight matrix and testing whether those neighborhood relationships are directionally preserved. The framework can therefore be applied to relationships between individual variables, between a focal taxon and the remainder of a community, or potentially between higher-dimensional groups of variables. Its most appropriate role is likely to be exploratory: identifying candidate directional relationships, prioritizing interactions for more detailed analysis, and generating hypotheses about influential components of ecological systems, particularly in large ecological datasets.

Several limitations define the current scope of the method. First, the simulation analysis was based primarily on a unidirectionally coupled logistic-map system. Although this system allowed causal direction and coupling strength to be controlled precisely, additional evaluation is needed for larger networks, bidirectional interactions, indirect causal pathways, common drivers, process noise, non-stationary dynamics, and irregularly sampled time series. In particular, the potential of FORCC for high-dimensional inference should be tested using simulated networks with known multivariate causal structures. Second, the quality of the forest-weight representation may depend on time-series length, embedding dimension, forest hyperparameters, and the minimum number of positive-weight neighbors. Sensitivity analyses across these choices will help identify practical guidelines for ecological applications. Third, the present implementation focused on near-immediate causal effects. Many ecological interactions operate with delayed or distributed effects, making systematic lag selection an important extension.

Fourth, surrogate-based inference depends on whether the chosen null model adequately represents the temporal structure of the data. Phase randomization preserves the approximate power spectrum and linear autocorrelation of an individual series while disrupting its phase relationships with other variables (Ebisuzaki, 1997). However, it may not fully reproduce non-stationarity, episodic environmental forcing, nonlinear seasonality, or dependencies created by unmeasured common drivers. Significant FORCC relationships should therefore not automatically be interpreted as direct biological interactions. They may also arise from indirect pathways or environmental variables not included in the analysis. Alternative surrogate procedures and conditional or multivariate extensions of FORCC may help distinguish these possibilities. Finally, broader comparisons with information-theoretic, graphical, and model-based causal-inference methods will be necessary to define the systems and data conditions under which FORCC provides the greatest benefit.

In conclusion, FORCC offers a new framework for causal inference from ecological time series by combining adaptive forest-weight neighborhoods with rank-based measures of directional dependence. The method recovered known causal direction in nonlinear simulations, reduced false-positive detections caused by strong synchronization, and identified interpretable asymmetries in a long-term marine community. Its principal contribution is not simply the replacement of one distance measure with another, but a shift from reconstructing causal-variable values through cross mapping to evaluating directional information directly from learned neighborhood structure. As ecological monitoring produces increasingly complex multivariate datasets, FORCC provides a practical approach for screening causal relationships, identifying candidate driver-like components, and generating testable hypotheses about the organization of natural systems.

## DATA AND CODE AVAILABILITY

The simulation datasets and experimental time series, except for the fish community monitoring data in Maizuru Bay (https://doi.org/10.5281/zenodo.1039387) (Ushio et al., 2018), are available from the corresponding author upon reasonable request. The underlying R scripts and implementation algorithms are subject to institutional intellectual property management related to a pending patent application (Japanese Patent Application No. 2026–87255) and will be made available upon reasonable request for non-commercial academic research and reproducibility verification purposes.

## ACKNOWLEDGEMENTS

I thank Dr. Takehiko Yamanaka for invaluable discussions and comments on the previous draft. A part of this research was conducted using the AI research supercomputer ‘SHIHOU’ at the National Agriculture and Food Research Organization (NARO).

## FUNDING INFORMATION

This study was supported by the following funds, JSPS KAKENHI Grant JP23751015 and JSPS KAKENHI Grant JP26K09087.

## COMPETING INTERESTS

The author declares a competing financial interest in the form of a pending patent application in Japan related to forest-weighted prediction and causal-inference methods presented in this study (Japanese Patent Application No. 2026–87255, titled “Prediction method, causal inference method, prediction apparatus, causal inference apparatus, and program”, by S.K.).

## AUTHOR CONTRIBUTIONS

Shigeki Kishi conceived the study, developed the methodology, performed the research and visualization, and wrote and edited the manuscript.

## Notes

https://doi.org/10.5281/zenodo.1039387

## REFERENCES

Aggarwal, C. C., Hinneburg, A., & Keim, D. A. (2001). On the surprising behavior of distance metrics in high dimensional space. In Database Theory — ICDT 2001 (pp. 420–434). Springer Berlin Heidelberg.

Athey, S., Tibshirani, J., & Wager, S. (2019). Generalized random forests. Annals of Statistics, 47(2), 1148–1178.

Chang, C.-W., Miki, T., Ushio, M., Ke, P.-J., Lu, H.-P., Shiah, F.-K., & Hsieh, C.-H. (2021). Reconstructing large interaction networks from empirical time series data. Ecology Letters, 24(12), 2763–2774.

Chatterjee, S. (2021). A new coefficient of correlation. Journal of the American Statistical Association, 116(536), 2009–2022.

Cheng, C., Sa-Ngasoongsong, A., Beyca, O., Le, T., Yang, H., Kong, Z. (james), & Bukkapatnam, S. T. S. (2015). Time series forecasting for nonlinear and non-stationary processes: a review and comparative study. IIE Transactions: Industrial Engineering Research & Development, 47(10), 1053–1071.

Chicharro, D., & Andrzejak, R. G. (2009). Reliable detection of directional couplings using rank statistics. Physical Review. E, Statistical, Nonlinear, and Soft Matter Physics, 80(2 Pt 2), 026217.

Cobey, S., & Baskerville, E. B. (2016). Limits to causal inference with state-space reconstruction for infectious disease. PloS One, 11(12), e0169050.

Ebisuzaki, W. (1997). A method to estimate the statistical significance of a correlation when the data are serially correlated. Journal of Climate, 10(9), 2147–2153.

Lin, Y., & Jeon, Y. (2006). Random forests and adaptive nearest neighbors. Journal of the American Statistical Association, 101(474), 578–590.

Mayr, M., Chasparis, G. C., & Küng, J. (2025). Causal time-series synchronization for multi-dimensional forecasting. Procedia Computer Science, 253, 2655–2664.

Misra, N. N., Dixit, Y., Al-Mallahi, A., Bhullar, M. S., Upadhyay, R., & Martynenko, A. (2022). IoT, big data, and artificial intelligence in agriculture and food industry.IEEE Internet of Things Journal, 9(9), 6305–6324.

Park, S. H., Ha, S., & Kim, J. K. (2023). A general model-based causal inference method overcomes the curse of synchrony and indirect effect. Nature Communications, 14(1), 4287.

R Core Team. (2025). R: A language and environment for statistical computing. R Foundation for Statistical Computing, Vienna, Austria. https://www.R-project.org/

Rulkov, N. F., Sushchik, M. M., Tsimring, L. S., & Abarbanel, H. D. (1995).Generalized synchronization of chaos in directionally coupled chaotic systems. Physical Review. E, Statistical Physics, Plasmas, Fluids, and Related Interdisciplinary Topics, 51(2), 980–994.

Runge, J., Bathiany, S., Bollt, E., Camps-Valls, G., Coumou, D., Deyle, E., Glymour, C., Kretschmer, M., Mahecha, M. D., Muñoz-Marí, J., van Nes, E. H., Peters, J., Quax, R., Reichstein, M., Scheffer, M., Schölkopf, B., Spirtes, P., Sugihara, G., Sun, J., … Zscheischler, J. (2019). Inferring causation from time series in Earth system sciences. Nature Communications, 10(1), 2553.

Runge, J., Gerhardus, A., Varando, G., Eyring, V., & Camps-Valls, G. (2023). Causal inference for time series. Nature Reviews. Earth & Environment, 4(7), 487–505.

Sugihara, G., May, R., Ye, H., Hsieh, C.-H., Deyle, E., Fogarty, M., & Munch, S. (2012). Detecting causality in complex ecosystems. Science (New York, N.Y.), 338(6106), 496–500.

Takens, F. (1981). Detecting strange attractors in turbulence. In Lecture Notes in Mathematics (pp. 366–381). Springer Berlin Heidelberg.

Tibshirani, J., Athey, S., Friedberg, R., Hadad, V., Hirshberg, D., Miner, L., Sverdrup, E., Wager, S., & Wright, M. (2026). Generalized random forests. CRAN.

Ushio, M., Hsieh, C.-H., Masuda, R., Deyle, E. R., Ye, H., Chang, C.-W., Sugihara, G., & Kondoh, M. (2018). Fluctuating interaction network and time-varying stability of a natural fish community. Nature, 554(7692), 360–363.

Wang, Y., Fang, L., & Jing, B. (2024). Adapted Chatterjee correlation coefficient.Statistics & Probability Letters, 215(110241), 110241.

Ye, H., & Sugihara, G. (2016). Information leverage in interconnected ecosystems: Overcoming the curse of dimensionality. Science (New York, N.Y.), 353(6302), 922–925.

